# The beta component of gamma-band auditory steady-state responses in patients with schizophrenia

**DOI:** 10.1101/2021.02.01.429120

**Authors:** Christoph Metzner, Volker Steuber

## Abstract

The mechanisms underlying circuit dysfunctions in schizophrenia (SCZ) remain poorly understood. Auditory steady-state responses (ASSRs), especially in the gamma and beta band, have been suggested as a potential biomarker for SCZ. While the reduction of 40Hz power for 40Hz drive has been well established and replicated in SCZ patients, studies are inconclusive when it comes to an increase in 20Hz power during 40Hz drive. There might be several factors explaining the inconsistencies, including differences in the sensitivity of the recording modality (EEG vs MEG), differences in stimuli (click-trains vs amplitude-modulated tones) and large differences in the amplitude of the stimuli. Here, we used a computational model of ASSR deficits in SCZ and explored the effect of three SCZ-associated microcircuit alterations: reduced GABA activity, increased GABA decay times and NMDA receptor hypofunction. We investigated the effect of input strength on gamma (40 Hz) and beta (20 Hz) band power during gamma ASSR stimulation and saw that the pronounced increase in beta power during gamma stimulation seen experimentally could only be reproduced in the model when GABA decay times were increased and only for a specific range of input strengths. More specifically, when the input was in this specific range, the rhythmic drive at 40Hz produced a strong 40Hz rhythm in the control network; however, in the ‘SCZ-like’ network, the prolonged inhibition led to a so-called ‘beat-skipping’, where the network would only strongly respond to every other input. This mechanism was responsible for the emergence of the pronounced 20Hz beta peak in the power spectrum. The other two microcircuit alterations were not able to produce a substantial 20 Hz component but they further narrowed the input strength range for which the network produced a beta component when combined with increased GABAergic decay times. Our finding that the beta component only existed for a specific range of input strengths might explain the seemingly inconsistent reporting in experimental studies and suggests that future ASSR studies should systematically explore different amplitudes of their stimuli. Furthermore, we provide a mechanistic link between a microcircuit alterations and an electrophysiological marker in schizophrenia and argue that more complex ASSR stimuli are needed to disentangle the nonlinear interactions of microcircuit alterations. The computational modelling approach put forward here is ideally suited to facilitate the development of such stimuli in a theory-based fashion.

## 1 Introduction

Auditory processing crucially relies on the fast temporal integration and resolution of inputs to form coherent percepts. Gamma band oscillations (>30 Hz) have been hypothesized to underlie this fast processing of auditory inputs [34, 33, 1] and, more generally, to establish communication between distributed neuronal groups [8]. One very simple way to test the ability of a neuronal microcircuit to generate and maintain oscillatory activity are steady-state responses (SSRs) - evoked oscillatory responses entrained to the frequency and phase of periodic stimuli. Importantly, patients with schizophrenia robustly show deficits in the 40-Hz auditory steady-state responses (ASSRs) [37, 21] and this general oscillatory deficit has been implicated in the pronounced perceptual and cognitive changes these patients experience [38]. This view is further underpinned by a large body of evidence documenting alterations of parvalbumin-positive (PV^+^) *γ*-aminobutyric acid (GABA) interneurons and their N-methyl-D-aspartate (NMDA) receptors [9, 10, 19, 22].

Interestingly, 50% of PV^+^ neurons in the dorsolateral prefrontal cortex of SCZ patients have very low levels of the GAD67 isoform of glutamate decarboxylase [16]. This reduced expression of GAD67 mRNA has been demonstrated to decrease the GABA synthesis in cortical GABAergic neurons, which would then result in smaller amplitudes of inhibitory postsynaptic currents (IPSCs) [9]. Additionally, PV^+^ neurons show reduced levels of the plasma membrane GABA transporter GAT1 in SCZ patients [42]. A reduced concentration of GAT1 has been shown to increase the time GABA molecules reside at the receptor and thus increase IPSC durations [32].

Administration of NMDAR antagonists lead to the emergence of schizophrenia-like symptoms, such as hallucinations, delusions and thought disorder, in healthy subjects [18]. Based on these findings it has been hypothesized that the reduced inhibition found in SCZ might not be a consequence of the changes to PV^+^ neurons described above, but could be attributable to an NMDAR hypofunction. Dysfunction of NM-DARs in SCZ is supported by several lines of evidence [19]. Specifically, Carlen et al.

[7] found that targeted deletion of NMDARs from PV^+^ interneurons led to increased spontaneous gamma oscillations and a deficit in gamma induction. Interestingly, they could reproduce these results in an established circuit model [40] when they implemented NMDAR hypofunction as an overall decrease in interneuron excitability.

In this study, we used an established network model of ASSR deficits in SCZ [40, 26], where SCZ-like behaviour is produced by an increase in IPSC decay times, to examine the dependence of 40 Hz ASSRs on the strength or amplitude of the inputs. In our model we could only reproduce the emergence of 20 Hz component during 40 Hz stimulation seen experimentally if the input strength was in a narrow range. More specifically, very weak input to the network did not result in a pronounced oscillatory rhythm. When the input was in a specific range, the 40 Hz stimulation entrained a pure 40 Hz oscillation in the control network, whereas in the ‘SCZ-like’ network, the changed IPSC time course caused a so-called ‘beat-skipping’, where the network would only strongly respond to every other input. This resulted in significant increase in 20 Hz power. Ultimately, if the input became too strong, the increased IPSC decay time was insufficient to suppress the very strong 40 Hz input. This was reflected in a single peak at 40Hz in the power spectrum. We then extended the network model to include more cellular-level alterations such as reduced GABA levels and NMDAR hypofunction. We found that the addition of further alterations did not change the input strength dependence of the 20 Hz component but further limited the parameter range where the component would occur. Our finding that the beta component only existed for a specific range of input strengths might explain the seemingly inconsistent reporting in experimental studies and suggests that future ASSR studies should systematically explore different amplitudes of their stimuli.

In this study, we used an established network model of ASSR deficits in SCZ [40, 26], where SCZ-like behaviour is produced in the model by an increase in IPSC decay times, to examine the dependence of 40 Hz ASSRs on the strength or amplitude of the inputs. We found that the pronounced increase in 20 Hz power during 40 Hz stimulation seen experimentally could only be reproduced in the model for a specific range of input strengths. More specifically, if the input was too weak the network failed to produce a strong oscillatory rhythm. When the input was in the specific range, the rhythmic drive at 40 Hz produced a strong 40 Hz rhythm in the control network, however, in the ‘SCZ-like’ network, the prolonged inhibition led to a so-called ‘beat-skipping’, where the network would only strongly respond to every other input. This mechanism was responsible for the emergence of the pronounced 20 Hz beta peak in the power spectrum. However, if the input exceeded a certain strength value, the 20 Hz peak in the power spectrum disappeared again. In this case, prolonged inhibition due to the increased IPSC decay times was insufficient to suppress the now stronger gamma drive from the input, resulting in an absence of the beat-skipping and single peak at 40Hz in the power spectrum. We then extended the network model to include more cellular-level alterations such as reduced GABA levels and NMDAR hypofunction. We found that the addition of further alterations did not change the input strength dependence of the 20 Hz component but further limited the parameter range where the component would occur. Our finding that the beta component only existed for a specific range of input strengths might explain the seemingly inconsistent reporting in experimental studies and suggests that future ASSR studies should systematically explore different amplitudes of their stimuli.

## 2 Methods

The model proposed here is based on a recent reimplementation [26] of the simple model presented by Vierling-Claassen et al. [40], which has been used in previous studies of ASSR deficits [30], and which is integrated in the ASSRUnit model database, a framework for automated testing of ASSR models against observations from empirical studies [28].

### Single Cell Model

Single cells are represented as theta neurons (see e.g. [4] for an in-depth analysis and description of the theta neuron model).

The *k*th neuron in a network is described by the variable *θ*_*k*_, which represents the neuron state, and which is governed by the following equation

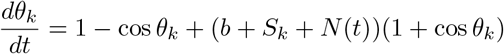

where *b* is an externally applied current, *S* is the total synaptic input to the cell and *N* (*t*) is a time-varying noise input. The total synaptic input to a cell in a network amounts to

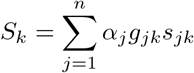

where *n* is the number of presynaptic neurons, *α*_*j*_ controls excitation and inhibition, i.e. is +1 for excitatory synapses and *-*1 for inhibitory ones, *g*_*jk*_ is the synaptic strength from cell *j* to cell *k* and *s*_*jk*_ is the synaptic gating variable from cell *j* to cell *k*. Synaptic gating variables evolve according to

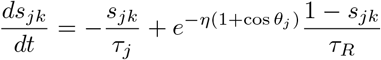

where *τ*_*j*_ is the synaptic decay time, *τ*_*R*_ the synaptic rise time and *η* is a scaling parameter. A single pacemaker cell provides rhythmic ASSR drive to the network. Additionally, Poissonian noise input is also given to all cells in the network, where a noise spike at time *t*_*n*_ elicits the following excitatory postsynaptic potential (EPSP) N(t)

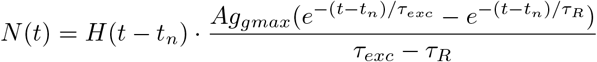

where *Ag*_*gmax*_ is the noise strength, *τ*_*exc*_ is the synaptic decay time, *τ*_*R*_ the synaptic rise time, and *H* the Heaviside function.

### Network

We combined 20 excitatory pyramidal cells together with 10 inhibitory cells into a network model, following the earlier work of [40, 26].

A schematic depiction of the network can be found in Figure 1. Populations connect to each other and also to themselves. The connectivity between any two populations is all-to-all. All populations also have two sources of input, the oscillatory drive input and a background noise input. The drive input periodically sends spikes with a given drive frequency to all populations, mimicking the rhythmic ASSR input. An overview of the model parameters can be found in Table 1.

**Figure 1:**
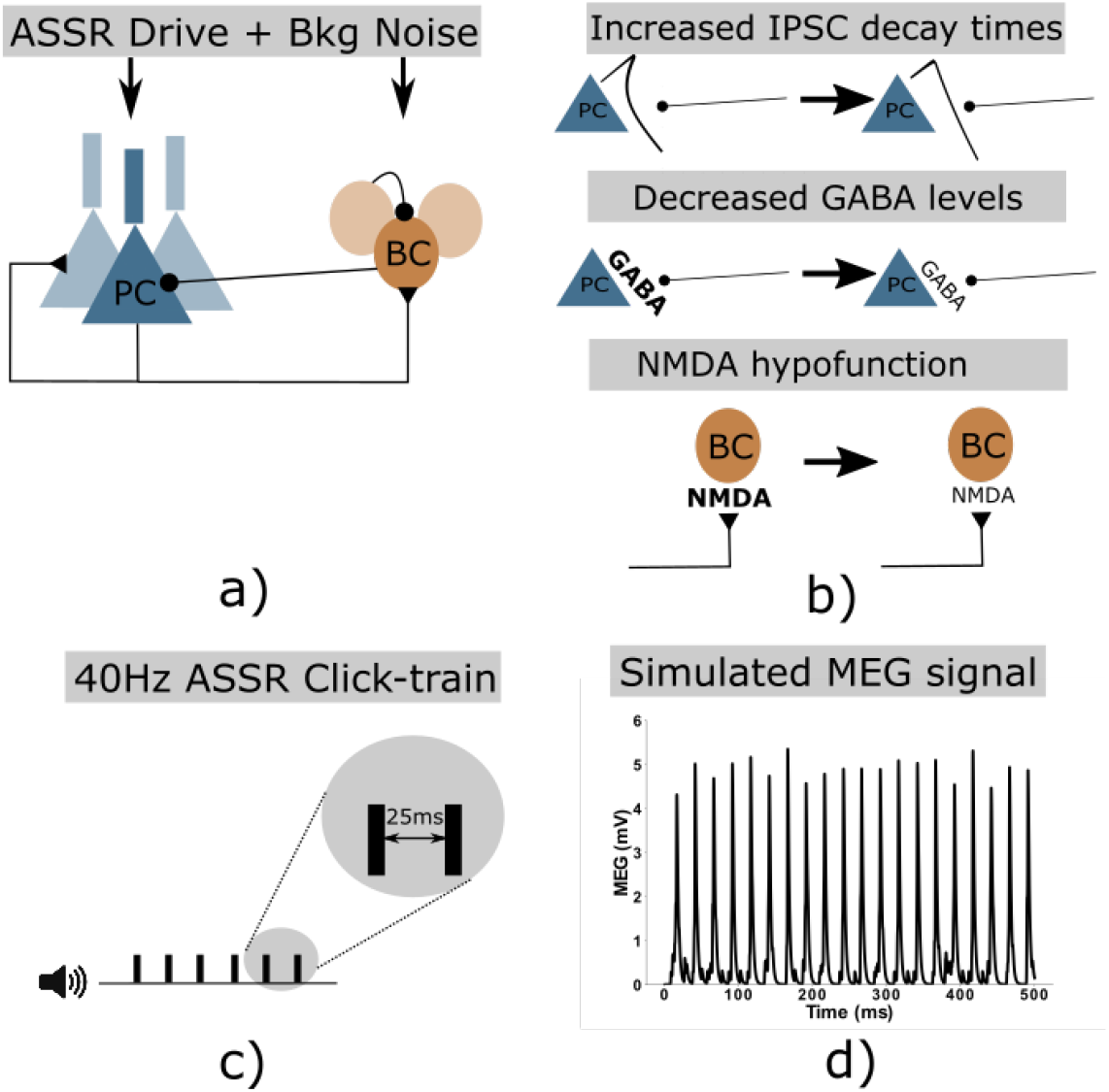
a) Network schematic showing the two neural populations (excitatory pyramidal cells and inhibitory basket cells) and their connectivity. Additionally, both populations receive periodic ASSR input drive and random background noise. b) Three potential microscopic changes underlying gamma ASSR deficits were implemented: Increased IPSC decay times at inhibitory synapses onto PCs, decreased GABA levels resulting in reduced IPSC amplitudes at inhibitory synapses onto PCs, NMDAR hypofunction resulting in decreased excitability of GABAergic interneurons c) Depiction of a 40 Hz click-train stimulus, where tones (synchronous inputs to the cells of the network) are presented with an inter-click interval of 25 ms resulting in a drive frequency of 40 Hz. d) Example simulated MEG signal of the network in response to a 40 Hz click-train stimulus.

**Table 1:** Model parameters

| Parameter | Definition | Value |
| --- | --- | --- |
| $n_E$ | Exc. population size | 20 |
| $n_I$ | Inh. population size | 10 |
| $\tau_R$ | Synaptic rise time | 0.1 |
| $\tau_{exc}$ | Excitatory decay time | 2.0 |
| $\tau_{inh}$ | Inhibitory decay time | 8.0 |
| $g_{ee}$ | E-E synaptic strength | 0.015 |
| $g_{ei}$ | E-I synaptic strength | 0.025 |
| $g_{ie}$ | I-E synaptic strength | 0.015 |
| $g_{ii}$ | I-I synaptic strength | 0.02 |
| $g_{de}$ | Synaptic strength of drive to E cells | 0.3 |
| $g_{di}$ | Synaptic strength of drive to I cells | 0.08 |
| $I$ | Input strength factor | varied |
| | (multiplied with $g_{de}$ and $g_{di}$ ) | (default=1.0) |
| $b$ | Applied current (regardless of cell type) | -0.1 |
| $Ag_{max}$ | Scaling factor for noise EPSCs | 0.6 |

To evaluate the oscillatory entrainment we calculate simulated EEG/MEG signals by summing all excitatory synaptic variables over all pyramidal cells (as in [40, 26, 30]). As the main measures for entrainment we perform a Fourier transform on these ‘EEG/MEG’ signals and extract the power at 40 Hz and at 20 Hz.

### Implementation of schizophrenogenic microcircuit alterations

We implemented changes to the GABAergic and glutamatergic neurotransmitter systems that have been associated with schizophrenia.

### GABAergic system

50% of PV^+^ neurons in the dorsolateral prefrontal cortex lack detectable levels of GAD67 [16]. It has been suggested that the reduced expression of GAD67 mRNA likely implies a reduction in GABA synthesis in cortical GABAergic neurons, which in turn would lead to smaller amplitude IPSCs at the postsynaptic site [9]. We implemented this change as a reduction of the weight of inhibitory connections (for both I-E and I-I connections).

Furthermore, a reduction in the plasma membrane GABA transporter GAT1 [32] has also been found in PV^+^ interneurons in SCZ patients [42]. GAT1 is a major contributor to the specificity of synapses by preventing spillover to neighbouring synapses [32] and a reduction in GAT1 results in prolonged IPSC durations [32]. Here, we realised this change as an increase of the IPSC decay time constant (as in previous studies [40, 26]).

### Glutamatergic system

NMDAR antagonists, such as phencyclidine and ketamine, produce symptoms, which are very similar to key clinical features of SCZ [18]. Convergent lines of evidence underpin that NMDARs are dysfunctional in SCZ [19]. Examples of direct evidence in favour of this hypothesis are changes in NMDAR-associated protein levels at the postsynaptic site [3], a reduction in NMDAR-mediated signalling following neuregulin 1 activation of ErbB4 receptors [11], lower levels of glutathione (a modulator at the redox-sensitive site of NMDARs) [36] and a reduction of kynurenine 3-monooxygenase that might increase kynurenic acid (an NMDAR antagonist) levels [41]. Indirect evidence, such as findings that putative risk genes for SCZ can affect NMDAR function [13] and that substances enhancing NMDARs might reduce symptom severity in SCZ [24], further underpin this idea. A potential hypofunction of NMDARs would lead to lower levels of excitation of PV^+^ interneurons and we therefore implemented this alterations by decreasing the applied current *b*_*inh*_ to inhibitory cells.

In this study, we considered four different ‘SCZ-like’ networks, which comprised the following combinations of changes to the GABAergic and glutamatergic system described above:

- *IPSC-SCZ-like* network: For this ‘SCZ-like’ network, we only implemented the increase of the IPSC decay time constant (as in previous studies [40, 26]).
- *IPSC+gGABA-SCZ-like* network: Here, additionally to the increase of IPSC decay times, we also reduced the weight of the inhibitory GABAergic connections (as in other previous studies [29]).
- *IPSC+bInh-SCZ-like* network: For this network, we decreased the applied current to the inhibitory cells together with the increase in IPSC decay times.
- *Full-SCZ-like* network: Here, we combined all three alterations, i.e. we implemented an increase in IPSC decay times, a decrease in inhibitory weights and a decrease in applied currents to inhibitory cells.

### Implementation details and code availability

The model was implemented using Python 2.7.9 and numpy 1.9.3. Analysis and visualization of the model output was also done in Python using the numpy and matplotlib packages (matplotlib 1.4.3).

Model equations were numerically solved using a simple forward Euler scheme. A single model run simulated a 500 ms trial and the time step was chosen such that this resulted in 2^13^ = 8192 data points. The model output was unaffected by using a smaller time step.

Simulation results varied from trial to trial because of the stochastic nature of the background input. Therefore, we always performed 20 simulation trials, each with a different realisation of the noise process. We then averaged these trials in time to get an average simulated MEG signal and all analyses were based on this average signal.

Model and analysis code are available on GitHub (https://github.com/ChristophMetzner/Gamma-Input-Dependence) and the model will be submitted to ModelDB (https://senselab.med.yale.edu/modeldb/) upon publication.

## 3 Results

### Replication of previous findings

First, we validated the *IPSC-SCZ-like* model against experimental observations [40] and replicated the findings from previous modelling studies with this model [40, 26]. The control network strongly entrains to the driving stimulus, regardless of the specific driving frequency (Figures 2 and 3 left columns), and shows stronger entrainment at 40 Hz than for 30 and 20 Hz, consistent with experiments [40]. Furthermore, the control model replicates another important feature seen in experiments and the previous models, a strong 40 Hz component for 20 Hz drive (Figures 2 and 3 left columns, third rows). The ‘SCZ-like’ network, where ‘SCZ-like’ behaviour is achieved by an increase in the GABAergic IPSC decay time constant (from 8 ms to 28 ms as in earlier studies), also reproduces important characteristics from experiments and previous models: First, the ‘IPSC-SCZ-like’ network shows a marked reduction in 40 Hz power for 40 Hz drive (Figure 3, right column, first row), as previously found in experiments (see [37] for a meta-analysis) and models [40, 26]. Furthermore, this network shows an emergent 20 Hz component at 40 Hz drive (Figure 3, right column, first row) as seen in [21, 40, 26] but not in other experimental studies [37]; and we see an increase in 20 Hz power and a relative decrease in 40 Hz power for 20 Hz drive in this condition (Figure 3, right column, third row).

**Figure 2:**
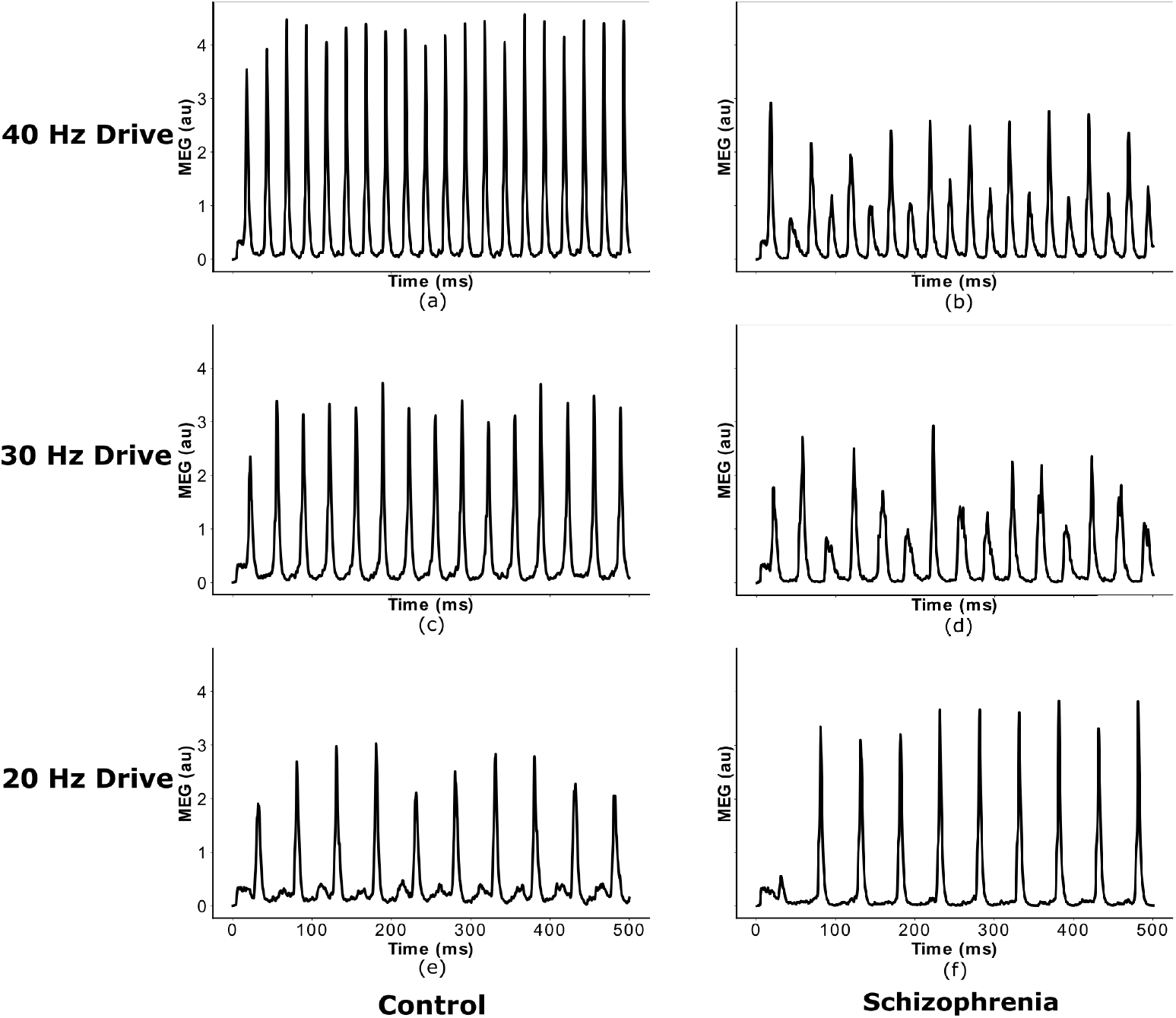
Network response to ASSR stimuli of different frequency. Simulated MEG signal of the control and *IPSC-SCZ-like* network in response to clicktrain stimuli with drive frequencies of 20, 30, and 40 Hz, replicating earlier studies using this model [40, 26].

**Figure 3:**
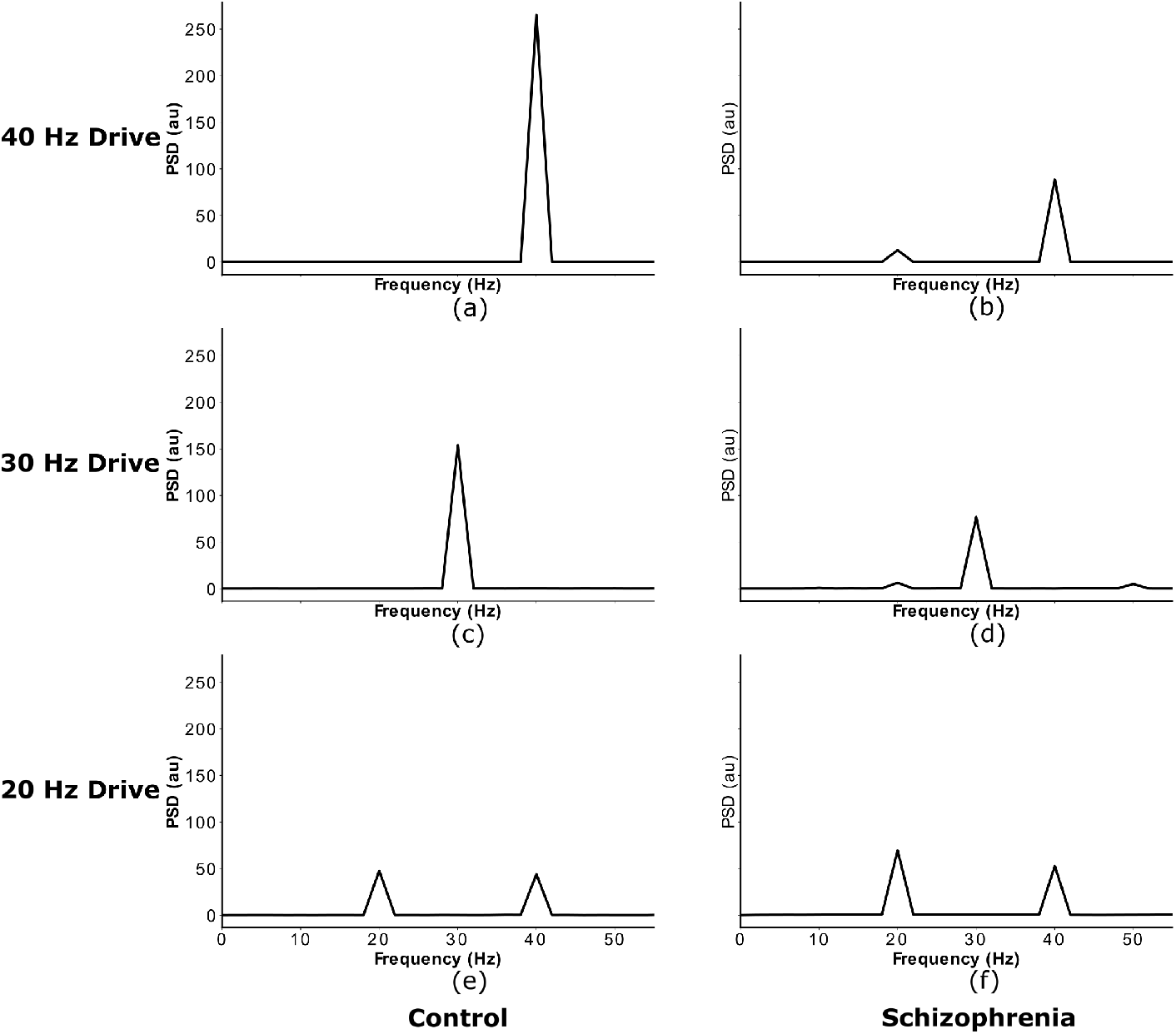
Power spectra of network responses to ASSR stimuli of different frequency. Power spectral densities of the simulated MEG signals from Figure 2, again replicating earlier studies using this model [40, 26].

### Input Strength Dependence of the 20 Hz Component

Next, we explored the input strength dependence of the *IPSC-SCZ-like* model response to 40 Hz drive by multiplying the input strength of the SCZ model by factors ranging from 0.1 to 1.5 in steps of 0.1. Figure 4 a) shows that the 40 Hz power increases with increasing input strength. Figure 4 b) shows that the 20 Hz component emerges at an input strength of 80% of the default strength, sharply increases for stronger inputs around the the default SCZ network strength and then sharply decreases again for inputs of 120% of the default strength and higher. Thus, in a narrow range between 80 and 120% of the default inhibitory input strength the network response exhibits a shift of power from the gamma (40 Hz) to the beta (20 Hz) band. Figures 4 c) and f) and Supplementary Figure 10 a) show that for weak inputs the oscillatory drive is not strong enough force the network into a coherent rhythm and, therefore, the powers at 40 Hz and at 20 Hz are very low. For input strengths around the default SCZ network value, the input strongly drives the network and forces it into a rhythm. However, the increased IPSC decay times prevent the excitatory pyramidal neurons from responding to every 40 Hz cycle and only allow them to spike every other cycle (Figures 4 d) and g) and Supplementary Figure 10 b)). Thus, the network rhythm displays a so-called ‘beat-skipping’ behaviour. The power spectrum of the response therefore shows both a 40 Hz (which is substantially smaller then for the control network) and a prominent 20 Hz peak (which is not visible for the control network). If the input, however, exceeds 120% of the default SCZ network value, the rhythmic input becomes strong enough to overcome the prolonged inhibition and forces the network into a gamma oscillation at 40 Hz. Here, the excitatory cells fire during each cycle and the power spectrum only shows a large 40 Hz peak (Figures 4 e) and h) and Supplementary Figure 10 c)).

**Figure 4:**
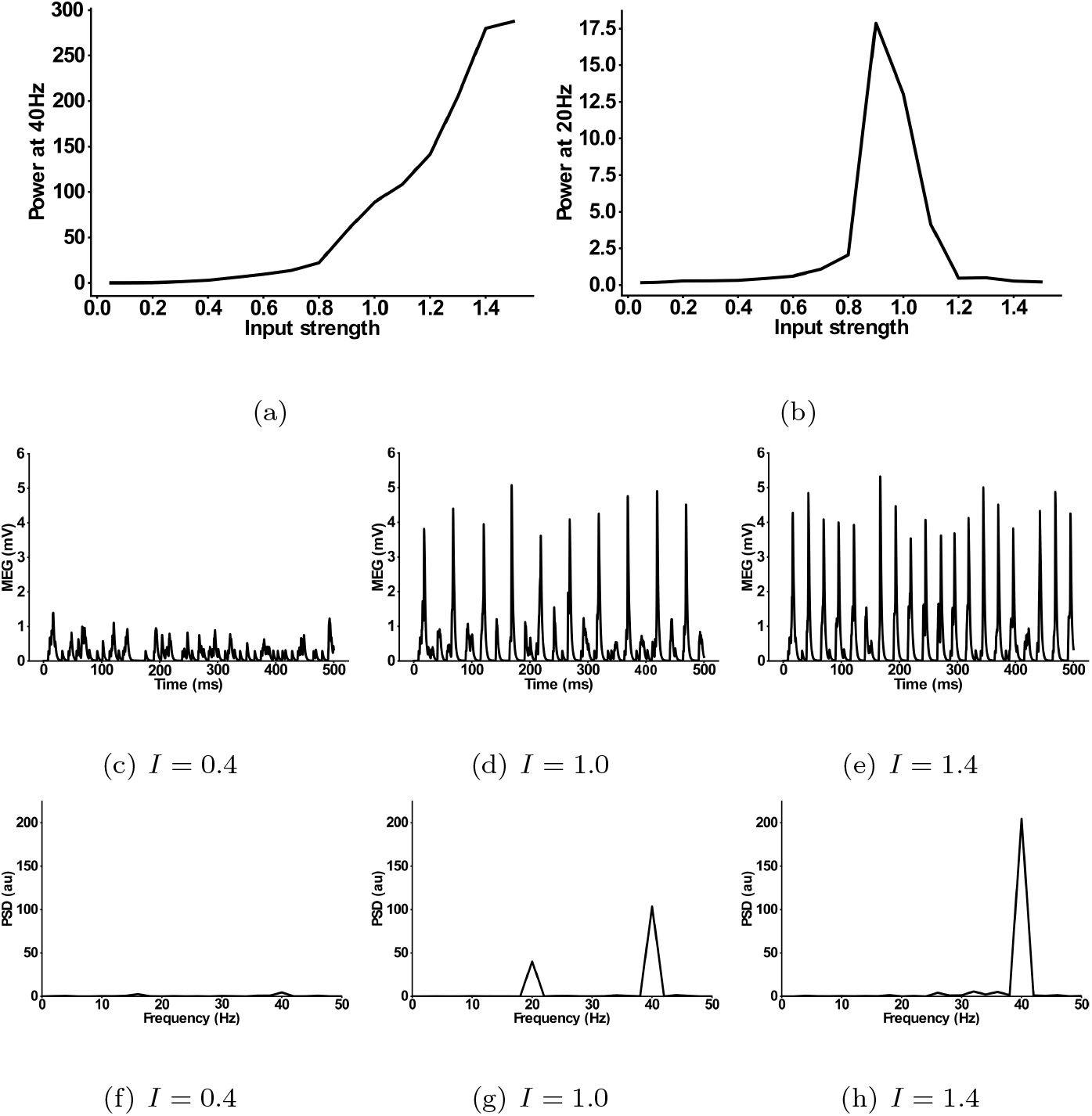
Input dependence of the 20 Hz component in the ‘IPSC-SCZ-like’ model. (a) Power at 40 Hz in response to 40 Hz drive as a function of the input strength. (b) Power at 20 Hz in response to 40 Hz drive as a function of the input strength. (c-e) Simulated MEG signals for three different input strengths: (c) *I* = 0.4 Input strength too low to drive synchronization. (d) *I* = 1.0 Input strength high enough to drive synchronization and to allow for a beat-skipping behaviour. (e) *I* = 1.4 Input strength too strong for beat-skipping behaviour, external 40 Hz drive dominates recurrent effects. (f-h) Power spectral densities for the signals from (c-e).

### Combinations of Alterations and their Input-Strength-Dependence

As explained earlier it is unlikely that the microscopic alterations associated with schizophrenia occur in isolation. Therefore, we added two more alterations to the *IPSC-SCZ-like* model: 1) A reduction in GABA levels, implemented as a reduction of the inhibitory weights; 2) A hypofunction of NMDARs at inhibitory interneurons, implemented as a reduction in interneuron excitability. We first added these modifications individually and combined them in a final set of simulations.

For the *IPSC+gGABA-SCZ-like* model, which included different GABA levels, we can see in Figure 5 that the 40 Hz component is shifted to lower input strengths and slightly decreased in power for low levels of GABA, but that the main effect is on the 20 Hz component. However, the emergent 20 Hz component, which existed for a narrow input strength range for the *IPSC-SCZ-like* model, narrowed down and finally vanished for stronger reductions of GABA levels. For the *IPSC+bInh-SCZ-like* model, with NMDAR hypofunction, the input strength dependence of the 40 Hz component exhibited a shift to lower input strengths than the model without NMDA receptor hypofunction and the 20 Hz component only emerged for weak reductions of the interneuron excitability (Figure 6). Lastly, the full model combining all three alterations, displayed similar behaviour as the previous models but an even more pronounced shift of the 40Hz components to lower input strengths for higher levels of GABA (Figure 7).

**Figure 5:**
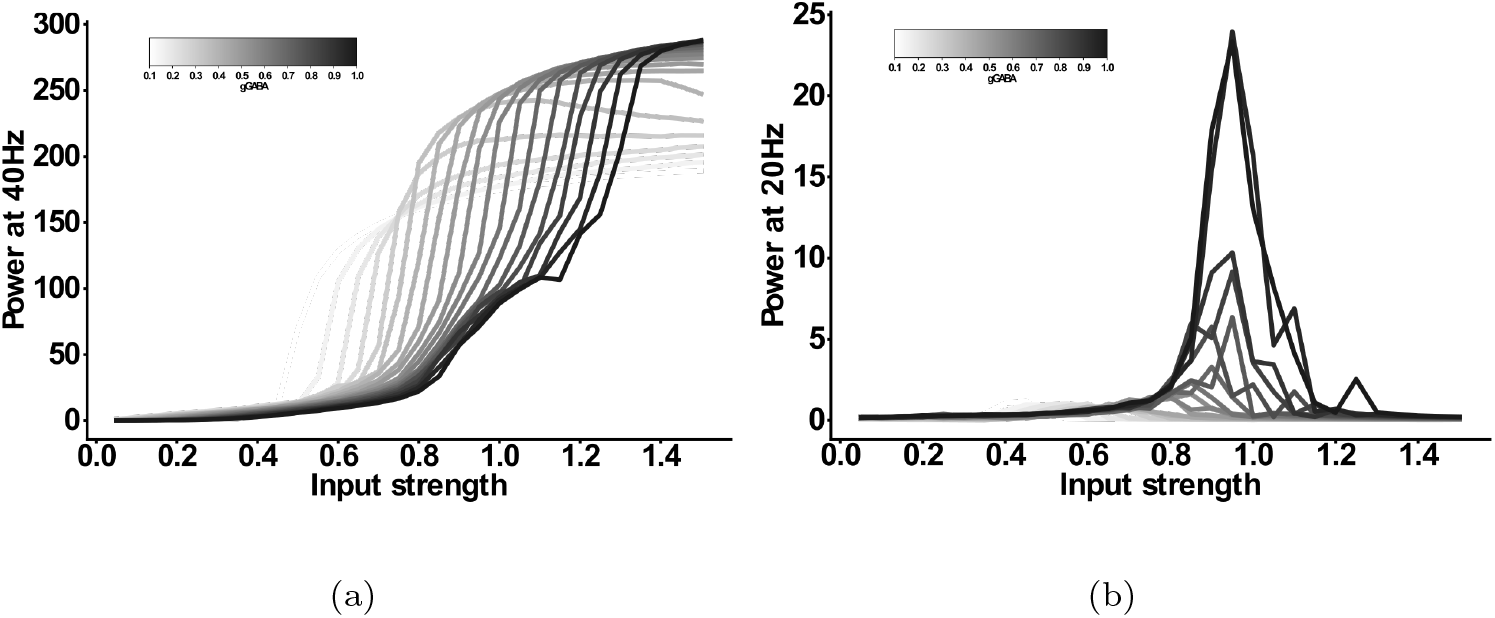
Input strength dependence of the 20 Hz component in the ‘IPSC+gGABA-SCZ-like’ model. (a) Power at 40 Hz in response to 40 Hz drive as a function of the input strength. (b) Power at 20 Hz in response to 40 Hz drive as a function of the input strength. In both plots the network model has increased IPSC decay times (from 8 ms to 28 ms) and the I-E and I-I synaptic strength (*g*_*ie*_ and *g*_*ii*_, respectively) is varied from 100% (black) to 10% (lightest grey) in steps of 5%.

**Figure 6:**
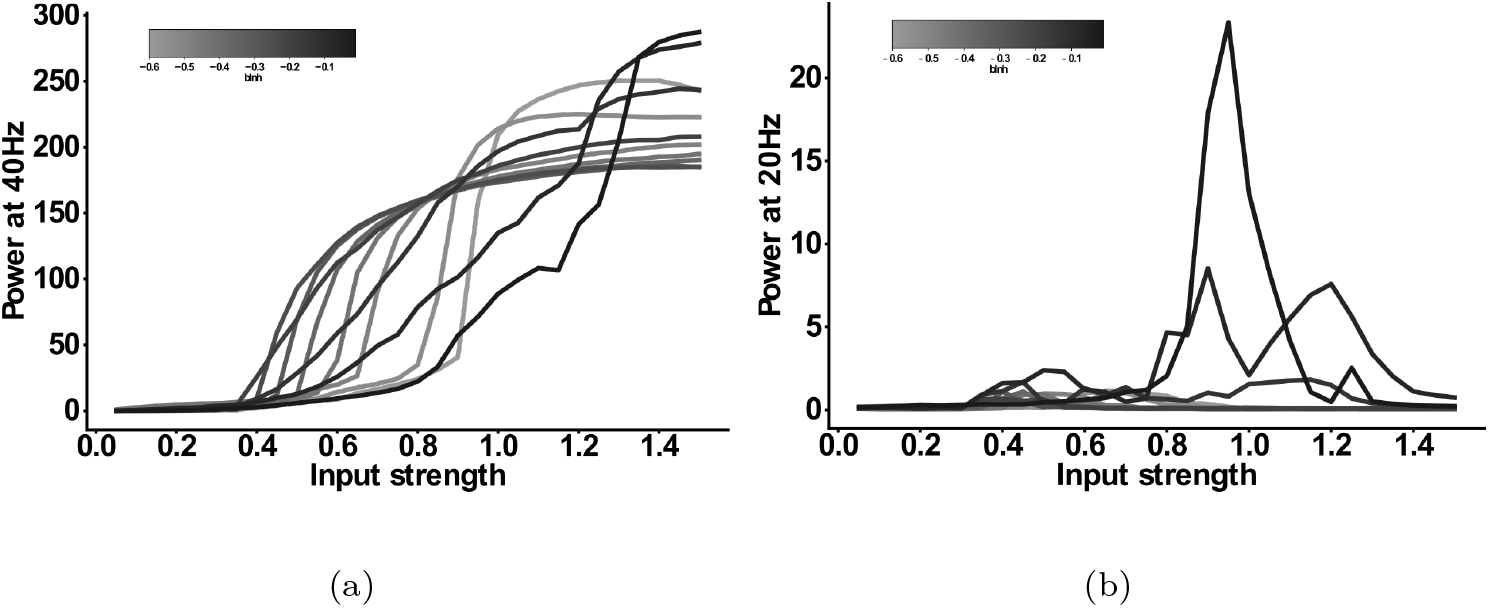
Input strength dependence of the 20 Hz component ‘IPSC+bInh-SCZ-like’ model. (a) Power at 40 Hz in response to 40 Hz drive as a function of the input strength. (b) Power at 20 Hz in response to 40 Hz drive as a function of the input strength. In both plots the network model has increased IPSC decay times (from 8 ms to 28 ms) and the interneuron excitability *b*_*inh*_ is varied from -0.01 (black) to -0.1 (darkest grey) and then in steps of -0.05 to -0.6 (lightest grey).

**Figure 7:**
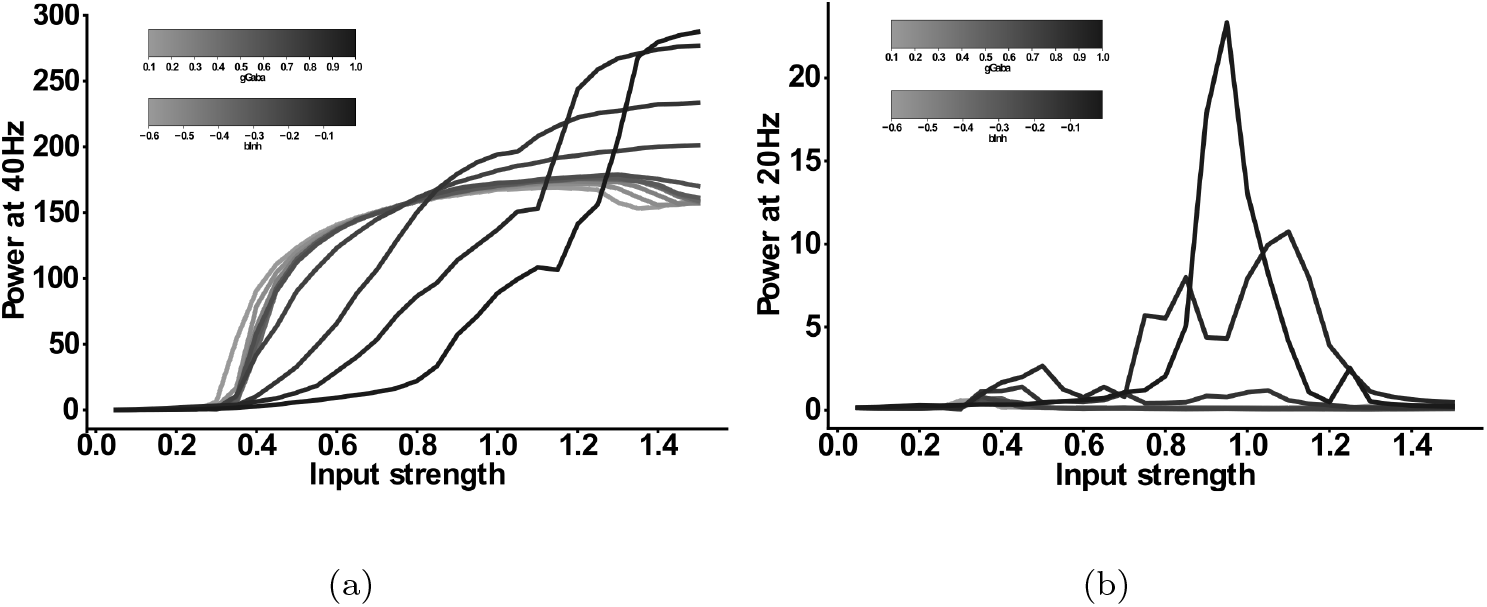
Input strength dependence of the 20 Hz component ‘Full-SCZ-like’ model. (a) Power at 40 Hz in response to 40 Hz drive as a function of the input strength. (b) Power at 20 Hz in response to 40 Hz drive as a function of the input strength. In both plots the network model has increased IPSC decay times (from 8 ms to 28 ms) and now both the I-E and I-I synaptic strength (*g*_*ie*_ and *g*_*ii*_, respectively) is varied from 100% (black) to 10% (lightest grey) in steps of 10% and simultaneously the interneuron excitability *b*_*inh*_ is varied from -0.01 (black) to -0.1 (darkest grey) and then in steps of -0.05 to -0.6 (lightest grey).

**Figure 8:**
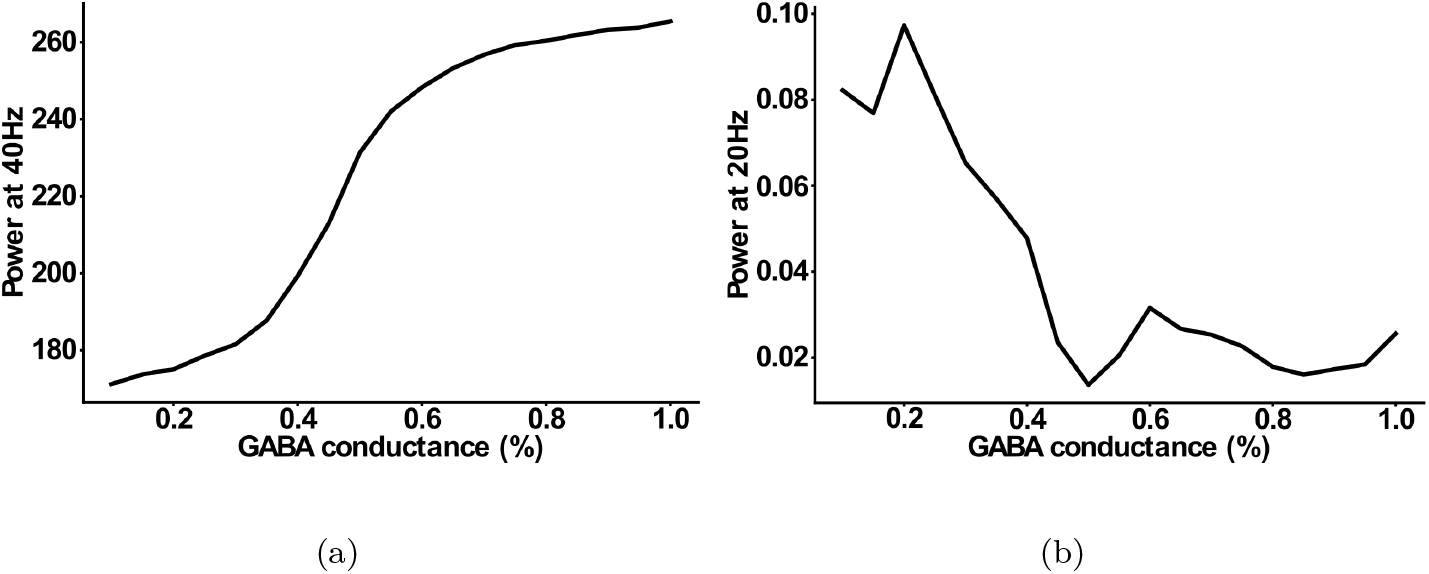
Effect of reduced GABAergic synaptic strength. (a) Power at 40 Hz in response to 40 Hz drive as a function of GABAergic synaptic strength. (b) Power at 20 Hz in response to 40 Hz drive as a function of GABAergic synaptic strength.

**Figure 9:**
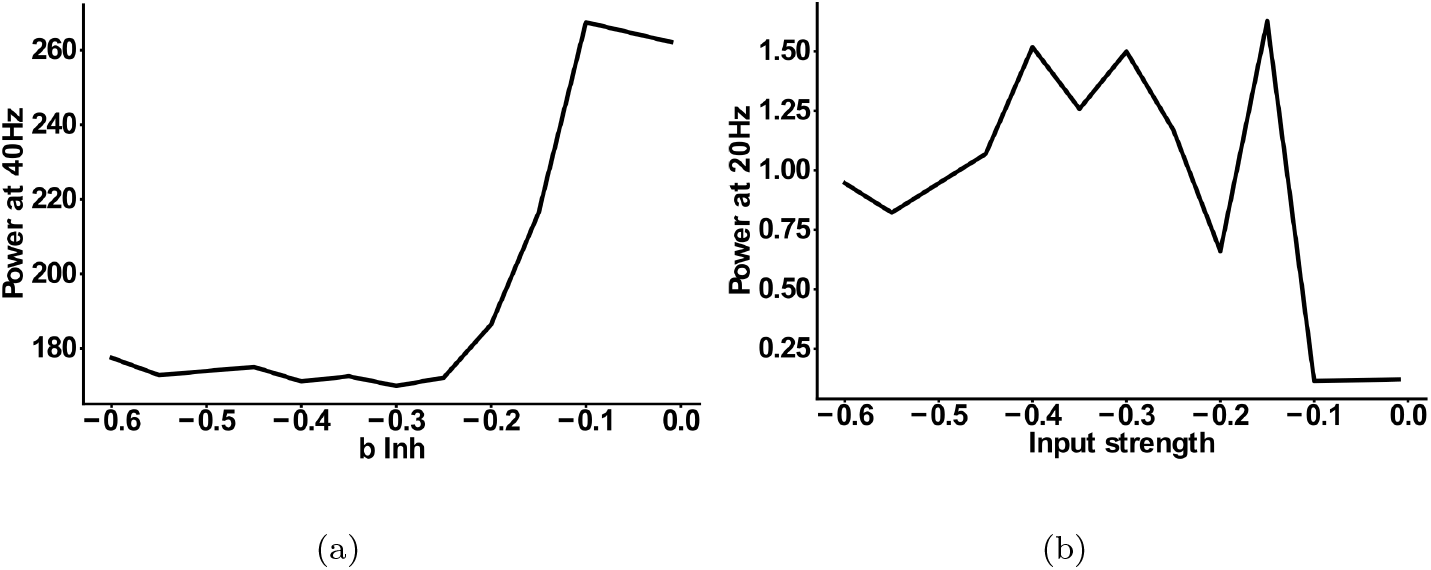
Effect of reduced interneuron excitability. (a) Power at 40 Hz in response to 40 Hz drive as a function of interneuron excitability. (b) Power at 20 Hz in response to 40 Hz drive as a function of interneuron excitability.

**Figure 10:**
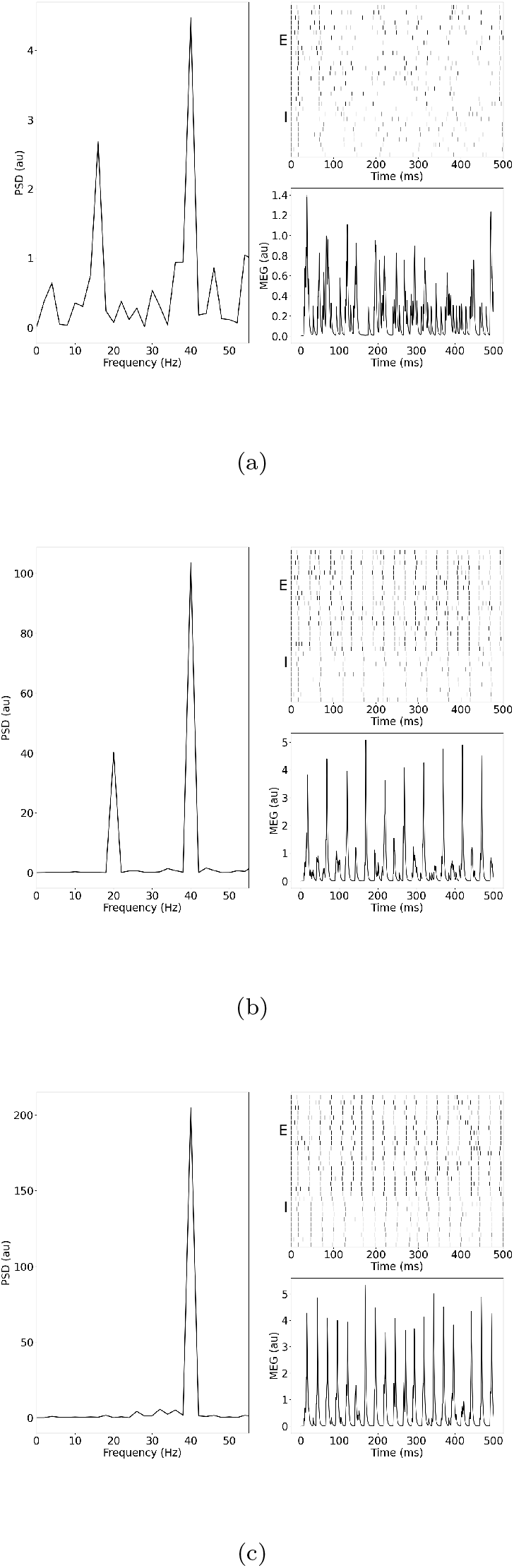
Single model trials. (a) Power spectral density, simulated MEG signal and raster plot for the IPSC model with an input strength of *I* = 0.4. (b) Same as (a) but for *I* = 1.0. (c) Same as (a) but for *I* = 1.4.

## 4 Discussion

Whether the presence of a 20 Hz beta response component in EEG/MEG signals for 40 Hz gamma drive during auditory steady state responses is an indicator of schizophrenia currently remains unresolved. In the present computational modelling study, we explored the conditions leading to the emergence of such a 20 Hz component. We could demonstrate that: (a) this beta component was only present in models that include an increased IPSC decay time but not in models that solely modelled decreased GABA activity or NMDAR hypofunction, further confirming the initial findings of Vierling-Claassen et al. [40], (b) the component strongly depended on the input strength and (c) the addition of GABA activity or NMDAR deficits to the IPSC decay time increases further narrowed the range of input strengths for which a substantial beta component existed. These results explain the seemingly inconsistent findings regarding the beta component of the 40 Hz ASSR measure in the literature.

However, there are several other potential factors that could contribute to these inconsistent experimental results. First, a difference in stimuli might play a role, since some studies use amplitude-modulated tones as opposed to the click-trains used in Vierling-Claassen et al. [40]. Second, most ASSR studies are performed using EEG recordings and the lower sensitivity of EEG compared to MEG might explain why the more subtle effect of the beta component has not been detected with EEG. Furthermore, as already pointed out by Vierling-Claassen et al. [40], averaging in time before the transformation into the frequency domain can potentially reduce the 20 Hz component considerably since the beat-skipping behaviour can vary from trial to trial. The skipped beats can be the 1st, 3rd, 5th,… in one trial while being the 2nd, 4th, 6th,.. in another trial, and thus would cancel out when averaged over in time prior to the frequency transform. Nevertheless, our modelling results offer a plausible and simple explanation of the inconsistent experimental results and suggest a careful choice of the input strengths in ASSR experiments.

We used a simple computational model consisting of an excitatory population, representing pyramidal cells, and an inhibitory population, representing PV^+^ inhibitory interneurons. While most of the experimental evidence for a reduction in GAT1, which in turn would lead to an increase in IPSC decay times, points towards chandelier cells [23], we have previously shown that at realistically low ratios of chandelier cells to basket cells in a microcircuit, gamma and beta range ASSR changes as seen in SCZ patients, are most likely due to an increase of IPSC decay times at basket cell synapses [30]. Our simplified model does not incorporate other types of inhibitory interneurons such as somatostatin-positive (SST^+^) or vasoactive intestinal peptide-positive (VIP^+^), although they have been shown to play important functional roles in cortical micro-circuits [5]. Furthermore, there is recent experimental evidence of alterations to SST^+^ interneurons in schizophrenia. Hashimoto et al. [14] found a reduced expression of GAD67 in SST^+^ neurons, while no reduction of GAT1 was apparent. Furthermore, Morris et al. [31] observed that both the density of SST neurons and the expression of SST^+^ per neuron was reduced in schizophrenia. These changes have been found in most cortical layers with varying strength [31] and can be observed throughout cortex [15]. This suggests, however, that IPSC decay times at SST^+^ interneuron synapses, a necessary requirement for the emergence of a beta component in our model, should remain intact in patients with schizophrenia. Additionally, the generation and maintenance of fast cortical rhythms in the beta and gamma range has been mainly attributed to PV^+^ neurons [12, 2, 6], although SST^+^ neurons have recently also been found to be involved [39]. These findings suggest that alterations of SST^+^ interneurons should only play a minor role in the emergence of a beta component in gamma ASSR tasks, and they were therefore not considered in the present study. Nevertheless, an exploration of the effects of SST^+^ alterations on cortical rhythms, especially for the lower frequency bands such as theta and alpha and for theta-gamma cross-frequency coupling, is warranted.

Beyond the question whether a beta component emerges in ASSR responses of patients with schizophrenia or not, our modelling work addresses a broader and more important issue. In general, it has proven extremely difficult to map schizophrenia-associated alterations of local microcircuits to specific neurocognitive or electrophysiological markers. Similar difficulties exist for other neuropsychiatric disorders such as autism spectrum disorder. Our work here shows that, while the robust deficit in the 40 Hz response to 40 Hz drive is not specific to a single microcircuit alteration, the emergence of the 20 Hz is. The computational model presented here mechanistically links the microcircuit change to the electrophysiological marker, thus, demonstrating the usefulness of mechanistic computational models in advancing our understanding of the relationship between features of the microcircuitry and non-invasive biomarkers, as we have argued before [25]. Furthermore, our simulation results show that the standard 40 Hz ASSR measure is not specific enough to resolve the complex, nonlinear interactions on the local circuit level and that more complex experimental designs are needed to disentangle them. This becomes especially important when considering that not only changes to the glutamatergic and GABAergic synapses considered in this work influence gamma ASSRs, but also neuromodulators such as dopamine [20] and cell-intrinsic changes to voltage-gated ionic channels [27]. This is further underpinned by the low specificity of the 40 Hz ASSR to schizophrenia, as for example similar ASSR deficits have been found in autism spectrum disorder [35] and bipolar disorder [17].

In summary, with this computational study we provide insights into the mechanistic generation of ASSR frequency components in schizophrenia beyond the traditional 40 Hz power at 40 Hz drive. Furthermore, we are able to explain seemingly conflicting experimental findings and suggest a more thorough and careful consideration of the effect of stimulus strength when designing ASSR experiments. Finally, we argue for a more complex and model-driven design of gamma and beta ASSR experiments in schizophrenia and for other neuropsychiatric disorders, which might be better suited to disentangle the nonlinear contributions of different microcircuit alterations found in these disorders.

## Supplementary Figures

### Reduced GABA levels

Here, we explore the effect of potentially reduced GABAergic conductances as a consequence of lower expression of GAD67 on the 40 and 20 Hz power at 40 Hz drive.

### NMDA hypofunction

Here, we explore the effect of potential NMDAR hypofunction on the 40 and 20 Hz power at 40 Hz drive.

### Input Dependence

Example single model trials for the IPSC model in three different regions of the input strength parameter space: below the critical range for the emergence of a 20 Hz component (10 (a)), within this range (10 (b)) and above (10 (c)).

